# Leveraging CRISPR activation for rapid assessment of gene editing products in human pluripotent stem cells

**DOI:** 10.1101/2024.09.29.615716

**Authors:** Youjun Wu, Aaron Zhong, Alessandro Evangelisti, Mega Sidharta, Lorenz Studer, Ting Zhou

## Abstract

Verification of genome editing in human pluripotent stem cells (hPSCs), particularly in silent locus is desirable but challenging because it often requires complex and time-intensive lineage-specific or tissue-specific differentiation to induce their expression. Here, we establish a rapid and effective workflow for the verification of hPSC lines with genome editing in unexpressed genes using CRISPR-mediated transcriptional activation (CRISPRa). We systematically compared the efficiency of various CRISPRa systems in hPSCs, identifying the SAM system as the most potent for activating silent genes in hPSCs. Furthermore, we demonstrated enhanced gene activation by combining the SAM system with TET1, a demethylation module. By inducing targeted gene activation in undifferentiated hPSCs using CRISPRa, we successfully verified single and dual reporter hPSC lines and conducted functional tests of dTAG knock-ins and silent gene knockouts within 48 hours. This approach eliminates the need for cell differentiation to access genes only expressed by differentiated cells, offering a handy assay for verifying gene editing in hPSCs.

## Introduction

CRISPR-Cas9 has been widely used for various of gene editing applications in hPSCs, targeting both expressed genes and silent genes in hPSCs^1,2^. Specifically, engineering of the silent genes in hPSCs such as lineage specific or tissue specific genes offers a robust platform for developmental studies, disease modeling, and drug screening^3,4^. Introducing an exogenous sequence into the endogenous locus or knocking out an endogenous gene are two critical and commonly used techniques in hPSC research. Knocking in a reporter gene or epitope tags into the endogenous locus provides a powerful tool for studying the biological function of genes of interest, isolating intended cell populations, investigating stem cell fate evolution and specification, and performing high-throughput screening^5–8^. Recently, knocking in conditional degron tags into the endogenous locus allows for tight control of proteins with small molecules, providing a powerful method to interrogate endogenous protein function by temporal dosage and reversible control^9–11^. Gene knockout by introducing a frameshift mutation or an early stop codon in the coding region of the target gene is the most the most straightforward approach for gene loss-of-function studies in hPSCs^12–14^.

Engineered knock-in or knockout hPSC lines are screened by genotyping with PCR and Sanger sequencing. However, evaluating the expression or function of the knock-in sequence and performing proteomic validation for the knockout gene remain the golden standards to confirm successful gene knock-in or complete depletion. Validation is not difficult when the target gene is expressed in parental hPSCs, but it becomes much more challenging when the targeted gene is not expressed in the parental cells. In such cases, verification requires differentiating hPSCs into a cell type that expresses the targeted genes. Some targeted genes only express in terminally differentiated cells, requiring a complicated, expensive, and time-consuming differentiation process. Moreover, the efficiency of the differentiation procedure directly affects verification, highlighting the urgent need for a more efficient and straightforward approach for verification of silent gene engineering in hPSCs.

Instead of activating silent genes by changing cell identity through differentiation, endogenous gene activation can be achieved through epigenetic modification of target genes. CRISPR activation (CRISPRa) technology uses catalytically inactivated Cas9 (dCas9) and sgRNA to recruit transcriptional activators, enabling gene activation from endogenous promoters^15,16^. This technology has been widely used for genetic screening, dissection of gene function in stem cell research, and as an alternative approach for human pluripotent reprogramming or cellular differentiation^17–20^. Given CRISPRa’s ability to induce expression of unexpressed genes, we explored its potential for routinely verifying hPSC lines with silent gene engineering.

In this study, we compared the robustness of the three most robust CRISPRa systems VPR, SAM and SPH and using a series of reporter hPSC lines targeting silent genes and identified SAM as the most potent for silent gene activation across all targets. We further found that incorporating an additional demethylation module, TET1, improves gene activation in highly methylated genes. We demonstrated the application of the CRISPRa system in the rapid verification of single or dual reporter gene activation and functional evaluation of dTAG knock-in and knockout hPSC lines as fast as 48 hours in undifferentiated hPSCs. Our study provides a systematic workflow for verifying hPSC lines with silent gene engineering without requiring complex cell differentiation which can be routinely applied to stem cell line generation.

## Results

### Comparison of various CRISPRa systems in hPSCs using lineage reporter lines

The three most potent CRISPRa systems, VPR^21^, SAM^22^ and Suntag-P65-HSF1 (SPH)^23^ were selected for evaluation across a series of reporter lines (Fig. 1A). These systems utilize different combinations of activation domains. The VPR system employs dCas9 fused with VP64, p65 and Rta (dCas9-VPR), alongside an sgRNA that directs the activators to specific targeted regions^21^. The SAM (synergistic activation mediator) system comprises dCas9 fused with VP64 (dCas9-VP64), a plasmid expressing two additional activators, P65 and HSF1, fused with the MS2 coated protein (MCP) (MCP-VP65-HSF1) and a modified sgRNA harboring MS2 stem loops (MS2-sgRNA), which recruit MCP-VP65-HSF1 and direct dCas9-VP64 to the target sites^22^. The SPH system, a more advanced version of Suntag system, uses multiple copies of GCN4 fused with dCas9 (dCas9-GCN4) to recruit transactivator P65 and HSF1, the two activator domains used in SAM, fused with a single chain variable fragment (scFv-P65-HSF1). This system was previously found more efficient than SAM and the original Suntag in HEK293T and N2a cells^23^ (Fig. 1A).

**Fig. 1:**
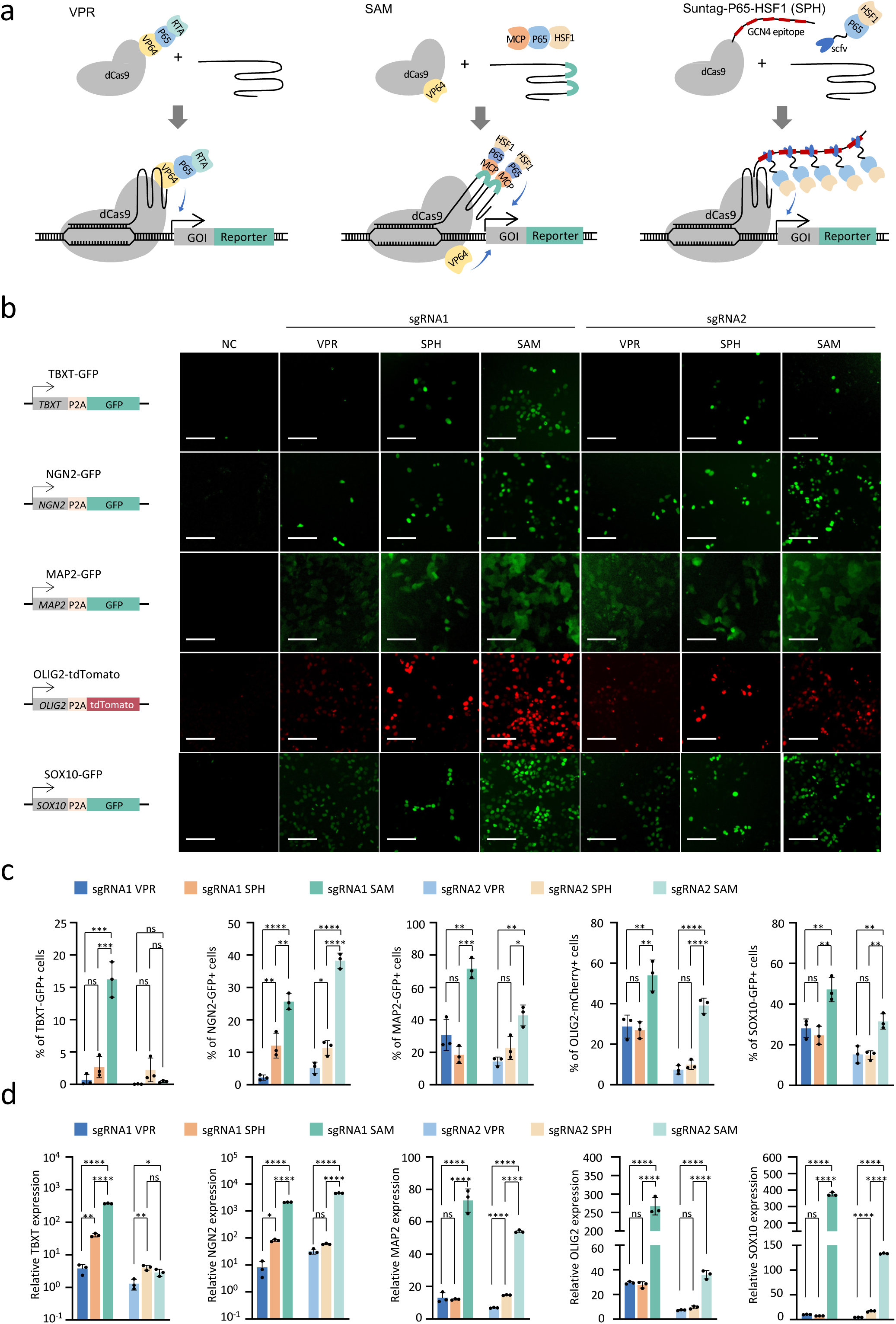
Comparison of different CRISPRa systems using hPSCs lineage reporter lines. **a.** Schematic of VPR, SAM and SPH CRISPRa systems. **b.** Fluorescence images of showing reporter gene activation with different CRISPRa systems in the hPSC lineage reporter lines. Two sgRNAs (sgRNA1 and sgRNA2) were tested for each lineage genes, including TBXT, NGN2, MAP2, OLIG2 and SOX10. Images were taken 48 hours post-electroporation. Each reporter cell line without electroporation was used as the negative control (NC). Scale bar: 10 µm. **c.** FACS analysis of fluorescence-positive cell population by VPR, SPH and SAM along with either sgRNA1 or sgRNA2. n=3 independent experiment. Data are presented as mean ± S.D. *p* values were calculated by one-way ANOVA with Tukey’s multiple comparison test (ns: non-significant, \**p*<0.05, \*\**p*<0.01, \*\*\**p*<0.001, \*\*\*\**p*<0.0001). **d.** Target gene expression induced by three CRISPRa systems measured by realtime qPCR relative to the parental cells. Data are presented as mean ± S.D. from 3 independent experiment. *p* values were calculated by one-way ANOVA with Tukey’s multiple comparison test (ns: non-significant, \**p*<0.05, \*\**p*<0.01, \*\*\*\**p*<0.0001).

Five lineage-reporter hPSC lines were employed to evaluate the effectiveness of the three CRISPRa systems including the reporters for primitive streak and mesodermal differentiation: TBXT-GFP^5^; the neuron specific transcription factor: NGN2-GFP; the mature neuron differentiation maker: MAP2-mCherry; the oligodendrocyte transcription factor: OLIG2-mCherry, and SOX10-GFP^24^, a transcription factor crucial for the development and maturation of glial cells (Supplementary Fig.1). These reporter lines were generated by knocking in a fluorescence gene into the specific gene locus which is silenced in undifferentiated hPSCs and only expressed when the cells undergo specific differentiation into a particular cell type. Flow cytometry and fluorescent microscopy analysis confirmed the fluorescence on these reporter lines remained undetectable in undifferentiated hPSCs (Fig. 1B). To activate each gene reporter, two sgRNAs (sgRNA1 and sgRNA2) were designed, targeting different regions within −300 to 0 bases from gene transcription star site (TSS). Each sgRNA was introduced along with other components of different CRISPRa systems through nucleofection in the reporter lines.

Fluorescence could be detected as early as 24h to 48 hours after nucleofection, although the potency of each CRISPRa system varied (Fig. 1B). Specifically, with sgRNA1, the TBXT-GFP signal was difficult to detect in cells treated with VPR 48 hours after nucleofection. In contrast, 2.7% of GFP-positive cells were observed with SPH, and over 16% with SAM. On the other hand, sgRNA2 did not perform well across all three systems, suggesting the targeting locus impacts CRISPRa efficiency. For NGN2-GFP activation, GFP was detectable with all three systems, showing 2.2%, 12%, and 25.6% GFP-positive cells when using sgRNA1; and 5.2%, 11.4%, and 38.3% GFP-positive cells when using sgRNA2, induced by VPR, SPH, and SAM respectively. In the cases of MAP2, OLIG2, and SOX10 activation, both VPR and SPH showed similar percentages of fluorescence-positive cells with either sgRNA1 or sgRNA2, while the SAM system significantly improved activation efficiency over the other two. The highest proportion of fluorescence-positive cells was detected when using SAM combined with sgRNA1, showing 71.7%, 54%, and 47.2% fluorescence in MAP2, OLIG2, and SOX10 reporter lines respectively (Fig. 1C, Supplementary Fig. 2). To verify the observations in the reporter lines as shown by FACS data, we measured the mRNA levels of the five targeted genes under each condition using qPCR. Consistently, conditions that yielded higher percentages of fluorescence-positive cells also showed higher levels of target gene expression (Fig. 1D).

It is worth noting that cytotoxicity was observed following nucleofection with the different CRISPRa systems. Based on the number of live cells 48 hours after nucleofection, targeting the five genes in hPSCs, SAM and VPR showed similar cell viability, while SPH demonstrated significant cytotoxicity, reducing the number of live cells by half compared to the other two systems (Supplementary Fig. 3). Overall, these data demonstrate that the SAM system is the most effective CRISPRa system compared to VPR and SPH, while also showing less toxicity in hPSCs.

### Improvement of silenced gene activation in hPSCs by incorporating a demethylating module

Using this approach, we further tested additional hPSC reporter lines and observed that the fluorescence signal in some lines remained weak, even with the SAM system. For instance, the activation of the KLF17 gene, a naïve pluripotent stem cell marker, was particularly weak (Fig. 2B). We hypothesized that these target genes might be heavily methylated in primed hPSCs, making them more challenging to activate solely through transactivators. To address this, we coupled demethylation with transcriptional activation by incorporating a demethylating module, the catalytic domain of TET1 (TET1-CD)^25^, into the SAM system, creating SAM-TET1 (Fig. 2A). A much stronger GFP signal was observed under fluorescence microscope in the cells treated with SAM-TET1 (Fig. 2B). FACS data revealed that the percentage of GFP-positive cells was similar between SAM and SAM-TET1, both outperforming VPR and SPH, consistent with the results for other genes tested (Fig. 2C, Supplementary Fig. 4A). However, the fluorescence intensity was dramatically increased with SAM-TET1, showing a 2.3-fold increase with sgRNA1 and a 3.6-fold increase with sgRNA2 compared to SAM alone (Fig. 2C). Consistently, mRNA levels were significantly elevated in the SAM-TET1 group compared to SAM (Fig. 2F).

**Fig. 2:**
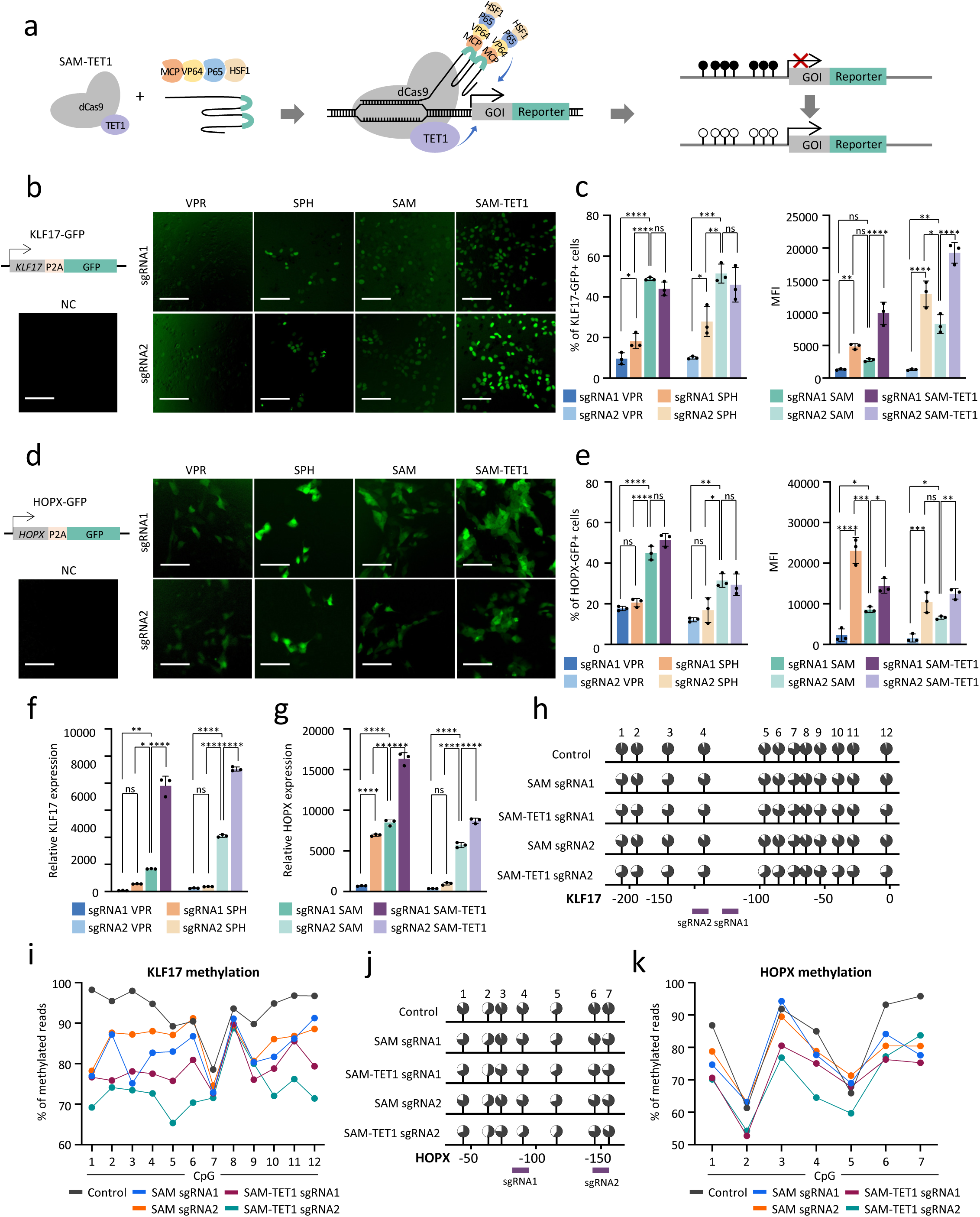
Combinatory effect of gene activation by SAM and demethylation module TET1CD. **a.** Schematic of generation of SAM-TET1 to activate genes silenced by hypermethylation. **b.** Reporter gene expression detected by fluorescence images in the H1-KLF17-GFP reporter cells 48 hours after electroporation with VPR, SPH, SAM or SAM-TET1 targeting KLF17 gene with two different sgRNAs. Scale bar: 10 µm. **c.** FACS analysis of the percentage of GFP-expressing cells (left panel) and median fluorescence intensity (MFI) of the GFP-positive cell population (right panel) in the reporter cells with KLF17 activation from b. **d, e.** Fluorescence signals upon HOPX activation in the H9-HOPX-GFP reporter cells by fluorescence images (d) and FACS (e), compared with VPR, SPH, SAM or SAM-TET1. **f, g.** KLF17 (f) and HOPX (g) gene expression measured by realtime qPCR activated the four systems. **h, i.** Methylation of region around sgRNA targeting sites of KLF17 gene in the undifferentiated hPSCs by SAM and SAM-TET1 along with two different sgRNAs, determined by targeted bisulfite sequencing. Cells electroporated with SAM along with empty sgRNA were set as the control. The distance of each CpG site to TSS, as well as the sgRNA target region, is showed on the x-axis (h). The percentages of methylated DNA at each CpG site are presented as black portion of the filled circles (h) or shown as individual dots in the line graph (i). **j, k.** Targeted bisulfite sequencing measuring the methylation percentage around sgRNA targeting region on the HOPX gene. Data in c, e, f and g are presented as mean ± S.D. from 3 independent experiment. *p* values were calculated by one-way ANOVA with Tukey’s multiple comparison test (ns: non-significant, \*\**p*<0.01, \*\**p*<0.001, \*\*\**p*<0.001, \*\*\*\**p*<0.0001).

We further compared SAM-TET1 derived activation with SAM, VPR and SPH in the HOPX-GFP reporter line^26^. Consistent with KLF17 gene activation, TET1 significantly improved reporter gene expression in the GFP-positive cell population when activating the HOPX gene (Fig. 2D). We found SPH produced the highest mean GFP intensity in this case, but the percentage of GFP-positive cells was lower than in both the SAM and SAM-TET1 groups (Fig. 2E, Supplementary Fig. 4B), which may be due to the toxicity of SPH in hPSCs (Supplementary Fig. 5). qPCR analysis showed that SAM-TET1 yielded the most potent transcriptional activation for HOPX (Fig. 2G). Meanwhile, unlike SPH, cell viability was not compromised with the addition of TET1 (Supplementary Fig. 5).

To further investigate, we assessed the methylation status of the KLF17 and HOPX target regions using targeted bisulfite sequencing in cells treated with SAM and SAM-TET1. The KLF17 target region in control cells with empty sgRNA was highly methylated, with 11 out of 12 CpG sites showing around or over 90% hypermethylation (Fig. 1H, I). Transcriptional activation of KLF17 by the SAM system led to a reduction in methylation compared to controls, reflecting the association between active transcription and lower DNA methylation levels (Fig. 1H, I). The addition of TET1 further decreased methylation at several CpG sites, particularly when combined with sgRNA2, where 10 out of 12 sites showed greater demethylation compared to SAM alone (Fig. 2H, I, Supplementary Fig. 6). In the case of HOPX activation, the initial methylation levels in control cells ranged from 61% to 95% across the 7 CpG sites. Transcriptional activation of HOPX resulted in reduced methylation at some sites, with TET1 inclusion further decreasing methylation compared to SAM alone (Fig.2J, K, Supplementary Fig. 6). This evaluation confirms that incorporating the demethylating module TET1 promotes demethylation the targeting locus, and further enhance the gene activation when coupled with CRISPRa-SAM.

### Rapid verification of reporter gene expression in unexpressed gene loci

The rapid and efficient activation of the fluorescence gene driven by silenced genes in hPSCs using CRISPRa suggests that this approach is a convenient and effective method for verifying hPSC reporter lines without requiring complex cell differentiation. Given that CRISPRa is a powerful tool capable of activating multiple genes simultaneously, we then applied it to verify dual reporter lines containing two different fluorescence genes inserted into two endogenous loci (Fig. 3A). Specifically, we generated a dual reporter hPSC line with GFP in frame with SOX10 and tdTomato in frame with OLIG2. This reporter line enables tracking changes in two critical transcription factors, SOX10 and OLIG2, during oligodendrocyte differentiation from hPSCs, and allow to isolate specific cell populations for mechanism studies. To verify the expression of the reporter genes, we introduced the SAM-TET1 system along with two efficient sgRNAs targeting SOX10 and OLIG2 promoters respectively. Both GFP and tdTomato were efficiently activated as early as 24-48 hours post-nucleofection (Fig. 3B). We also compared the activation efficiency of using two sgRNAs simultaneously versus using individual sgRNA targeting a single gene. FACS data showed that around 54.9% of cells were tdTomato-positive when targeting OLIG2 alone, and 52.4% GFP-positive when targeting SOX10 gene alone. Simultaneous targeting of both genes with two sgRNAs resulted in a similar population of reporter gene-positive cells, with 56.2 % expressing tdTomato, 51.4% expressing GFP, and 50.7% co-expressing both fluorescence genes (Fig. 3C).

**Fig. 3:**
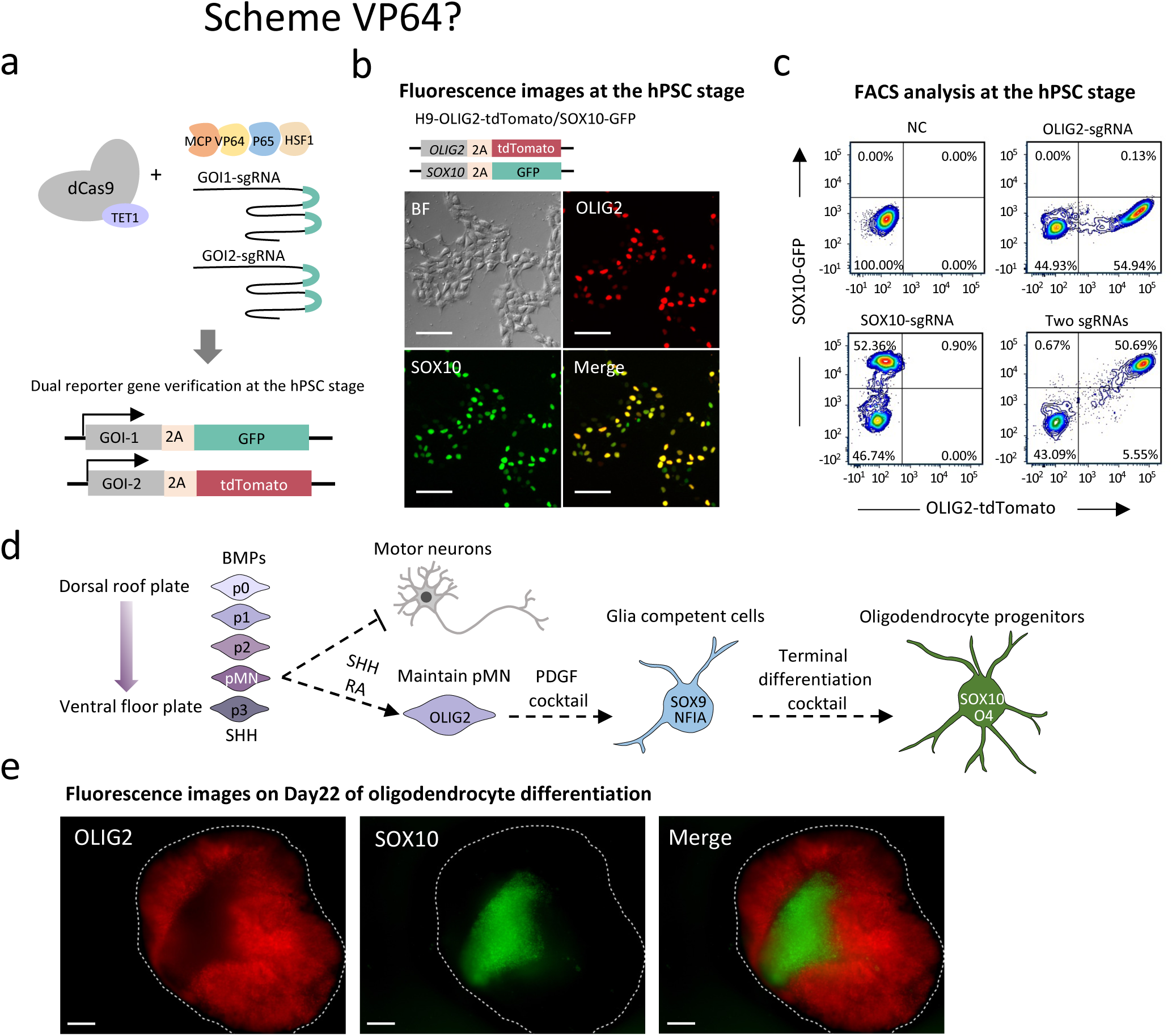
Verification of lineage dual reporter hPSC lines at the undifferentiated stage. **a.** Schematic of activating two reporter genes knocked into two unexpressed genes using CRISPRa. **b.** Fluorescence images of H9-OLIG2-tdTomato/SOX10-GFP cells 48 hours post-electroporation with SAM-TET1 and sgRNAs targeting OLIG2 and SOX10. tdTomato targeting the OLIG2 gene and GFP targeting SOX10, as well as the merged image, are demonstrated. BF, bright field. Scale bar: 10 µm. **c.** FACS analysis of tdTomato or GFP positive cells and the double-positive cell population 48 hours after electroporation. **d.** Schematic of the developmental process of oligodendrocytes differentiation. Oligodendrocytes originate from the pMN domain of the developing spinal cord where they express OLIG2. As they mature into glia-competent cells, they become locked into an oligodendrocytes competency state, with the help of factor such as PDGF, and become oligodendrocytes progenitor cells (OPCs) expressing nuclear SOX10. **e.** Fluorescence images of the differentiated cells on Day 22 of oligodendrocyte differentiation. The white dotted line outlines the organoid. Scale bar: 200 µm.

To determine whether the dual reporter cells verified by CRISPRa could be applied to hPSC differentiation, we conducted an oligodendrocyte differentiation using a stepwise protocol, where hPSCs were first differentiated into pMN cells expressing OLIG2 and subsequently transformed into oligodendrocyte progenitor cells (OPCs) expressing SOX10 (Fig. 3D). The dual fluorescence signals representing the two markers were detected around 20 days after differentiation. At this stage, the aggregates exhibited a large number of tdTomato-positive cells expressing OLIG2 and a proportion of GFP-positive cells expressing SOX10 (Fig. 3E). In summary, these data suggest that reporter hPSCs can be efficiently verified with CRISPRa at the undifferentiated stage, and importantly, the CRISPRa validated reporter hPSCs faithfully report the gene expression during the target lineage differentiation.

### Functional evaluation of knockdown of unexpressed genes targeted by degron tag in hPSCs

Degron tagging approach has been applied to control endogenous protein levels by knocking in degron tags at the endogenous locus^9–11^. We aimed to test whether CRISPRa could be used for the rapid functional evaluation of conditional degron tag systems knocked into the endogenous locus that is not expressed in hPSCs. In this study, we employed the dTAG system as an example, which utilizes the highly selective binding between an engineered FKBP^12F36V^ and the heterobifunctional dTAG molecule (dTAG-13)^10^. This interaction recruits the CRBN E3 ligase complex, inducing the degradation of the target protein (Fig. 4A).

**Fig. 4:**
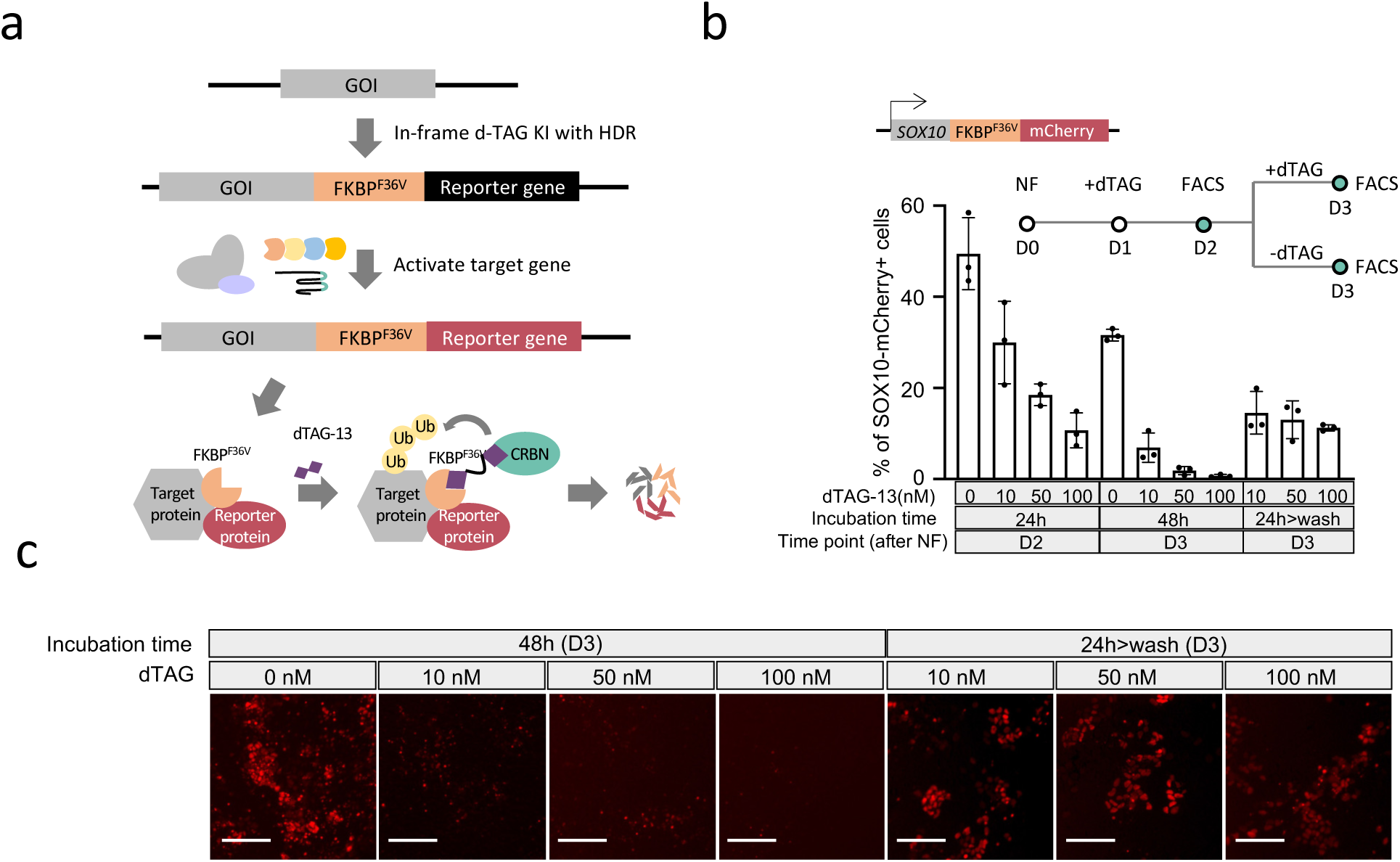
Functional evaluation of dTAG knocked in the silenced endogenous gene in undifferentiated hPSCs. **a.** Schematic of testing dTAG mediated conditional knockdown of targeted endogenous protein in undifferentiated hPSCs using CRISPRa. **b.** Evaluation of dTAG-mediated endogenous SOX10 degradation activated by CRISPRa. Upper panel: The experimental timeline for nucleofection with CRISPRa plasmids, dTAG-13 incubation, and washout, as well as the timepoints for analysis. Low panel: mCherry-positive cell population reflecting SOX10 expression was determined by FACS with the indicated treatment. **c.** Fluorescence images on Day 3 after nucleofection of the cells with 0-100 nM dTAG-13 for 48 hours or 24 hours followed by washout. Scale bar: 10 µm.

We introduced the FKBP12^F36V^-mCherry sequence into the C-terminal of the SOX10 gene in hPSCs through Cas9-mediated homology-directed repair (HDR) (Supplementary Fig. 7). This construct allows the fusion chimera SOX10-FKBP12F36V-mCherry to be induced for degradation with dTAG-13, and the expression of SOX10 can be easily monitored by checking the mCherry fluorescence.

To perform the functional assay of dTAG with the generated SOX10-FKBP^12F36V^-mCherry hPSC line, we first activated SOX10 expression using SAM-TET1. mCherry-positive cells were detected 24 hours post-nucleofection, indicating rapid SOX10 activation (Fig.4C). Different concentrations of dTAG-13, ranging from 0 to 100 nM, were added to the cells at this time point. Incubation with dTAG-13 resulted in a dose- and time-dependent decrease in the percentage of mCherry-positive cells (Fig. 4B, C). Specifically, around 50% of mCherry-positive cells were detected without dTAG upon SOX10 activation, which is comparable to the ratio of fluorescence-positive cells activated in the SOX10 reporter lines by SAM (Fig. 1C and 3C). Treatment with 10, 50, and 100 nM of dTAG resulted in 30.0%, 18.5%, and 10.7% of mCherry-expressing cells, respectively, after 24-hour incubation (Fig. 4B).

Since SOX10 was transiently activated by plasmid nucleofection, the mCherry-positive cell percentage decreased to 31.6% in the absence of dTAG-13 at 72 hours post-nucleofection. Treatment with dTAG-13 effectively ablated the majority of mCherry-positive cells, resulting in 6.9% with 10 nM dTAG-13 and less than 1% with higher concentrations (50 nM and 100 nM). These results suggest that the SOX10 protein induced by SAM was effectively degraded by dTAG-13 via the fused FKBP12F36V chimera.

Another important property of the dTAG platform is that the targeted protein degradation by the small molecule is reversible upon removal of the dTAG molecule. To assess this reversibility, we washed out dTAG-13 after 24 hours of incubation and evaluated the mCherry-positive cells again after another 24 hours. Upon washout, an increase in mCherry-positive cells was observed in all the cells treated with 10 nM, 50 nM, and 100 nM dTAG-13, showing 14.5%, 13%, and 11.3% mCherry-positive cells, respectively. Meanwhile, the no-drug control, consisting of cells post-nucleofection after 72 hours, demonstrated a 31.6% mCherry-positive population (Fig. 4B).

These results indicate that the SOX10 protein induced by SAM was effectively degraded by dTAG-13 and was partially recovered upon removal of the small molecule. In summary, all these data suggest that CRISPRa enables rapid evaluation of the function of the dTAG system, allowing conditional and reversible control of target endogenous proteins in undifferentiated hPSCs.

### Identification of knockout of unexpressed gene in hPSCs

Validating a gene knockout requires comparing the protein levels of the targeted gene between parental cells and knockout lines. However, when the targeted gene is silenced in hPSCs, we explored the possibility of using CRISPRa to identify successful knockout by inducing the expression of the target gene only in parental lines, not in knockout lines. To validate this approach, we first generated knockout lines for the SOX10 and KLF17 genes using CRISPR-mediated NHEJ, introducing out-of-frame mutations. We then measured the SOX10 and KLF17 proteins induced by SAM-TET1, which had previously demonstrated robust activation of these genes (Fig 2C and F, Fig. 3B and C).

For the SOX10 knockout in H1 cells, we isolated a single-cell clone that exhibited an “A” insertion in the coding region of SOX10 at the Cas9 cleavage site, leading to a frameshift mutation and an early “TGA” stop codon (Fig. 5B). After 48 hours of induction with SAM-TET1, SOX10 protein was highly expressed in the parental cells but was undetectable in this knockout clone under the same conditions (Fig. 5C), validated by western blot. Similarly, for the KLF17 knockout in the KLF17-GFP reporter line, Sanger sequencing revealed mutations around the Cas9 cleavage site, although the subsequent sequence was unreadable, an outcome sometimes observed when generating knockouts in hPSCs via Cas9-mediated NHEJ due to random frameshift mutations at both alleles (Fig. 5D). To further verify this cell line, we assessed the reporter gene in frame with KLF17 after SAM-TET1 treatment. In parental cells, 36% of cells were GFP-positive 48 hours after KLF17 activation, whereas GFP-positive cells were nearly undetectable in the knockout line (Fig. 5E).

**Fig. 5:**
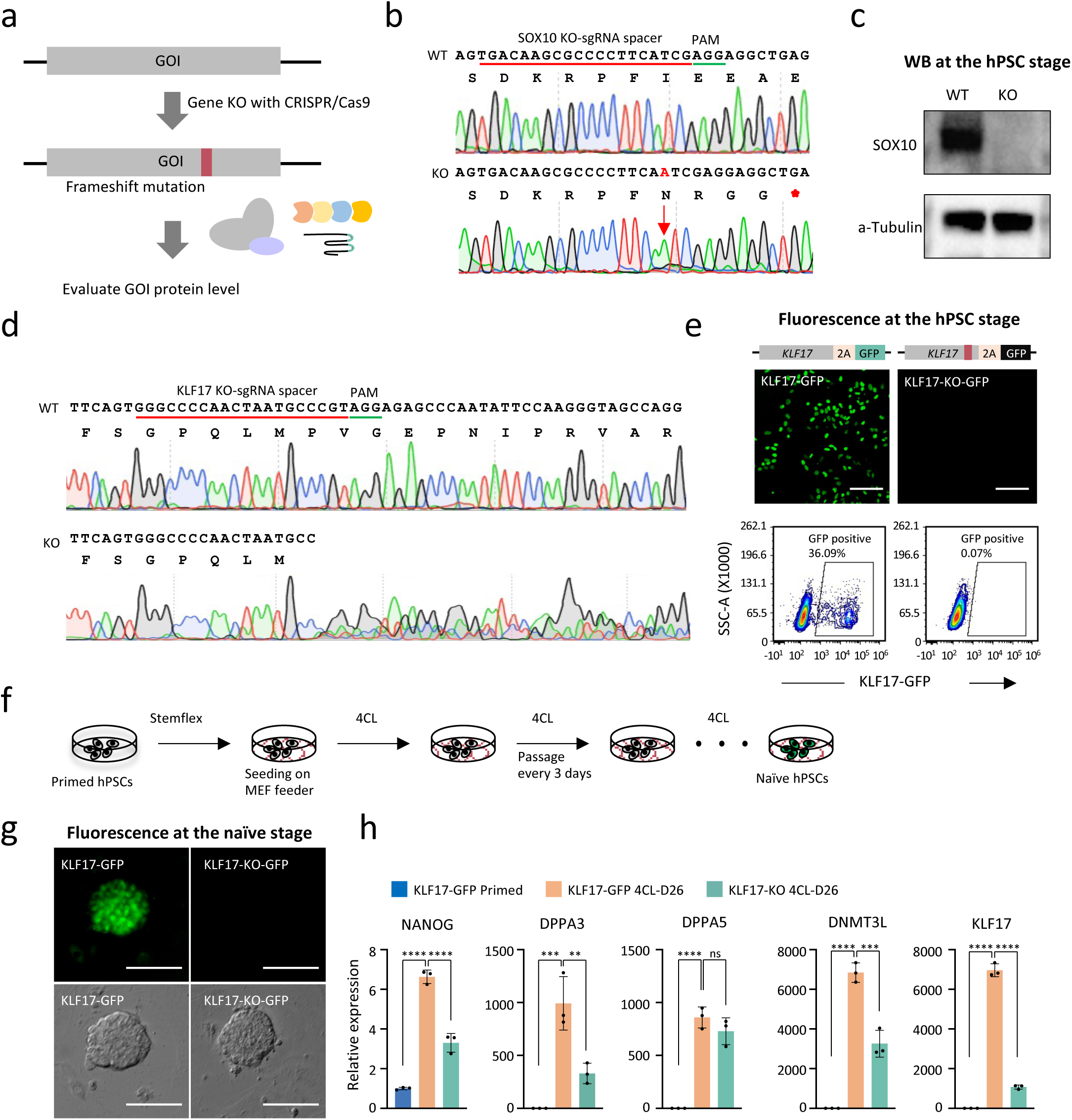
Verification of silent gene knockout in undifferentiated hPSCs using CRISPRa. **a.** Schematic of endogenous gene knockout using CRISPR/Cas9 and verification. **b.** Sanger sequencing of SOX10 gene in wild type H1 cell (WT) and single cell clone with CRISPRa/Cas9 Knockout targeting SOX10 (KO). The sgRNA targeting region (underline in red) and PAM (underline in green) are labeled. The KO line showed an “A insertion”, leading to a frameshift mutation and an early stop codon “TGA” labeled with asterisk. **c.** Western blot detection of SOX10 gene with SAM-TET1 activation in the WT and KO lines. **d.** Sanger sequencing of the KLF17 gene in the WT H1-KLF17-GFP cells and KLF17 KO cells. **e.** Fluorescence images (upper panel) and FACS analysis (lower panel) of GFP at the *KLF17* locus 48 hours with SAM-TET1 activation of KLF17 in both WT and KO cells. **f.** Schematic of naïve programming of hPSCs using 4CL medium. **g.** Fluorescence image detection of GFP expression in the H1-KLF17-GFP cells and the KLF17 KO cells on the Day 26 of naïve programming. Corresponding bright filed images of the naïve cells are showed in the lower panel. Scale bar: 10 µm. **h.** Expression of naïve hPSC-related genes in the parental and KLF17 KO cells using realtime qPCR. Data are presented as mean ± S.D. from 3 independent experiment. *p* values were calculated by one-way ANOVA with Tukey’s multiple comparison test (ns: non-significant, \*\**p*<0.01, \*\*\**p*<0.001, \*\*\*\**p*<0.0001).

KLF17 is not expressed in the primed state of hPSCs but is expressed in the naïve state^27,28^. To further confirm the results obtained with CRISPRa, we compared the KLF17-GFP reporter cells and the knockout line during naïve cell programming using the 4CL medium which effectively reprograms hPSCs from a primed to a naïve state^29^ (Fig.5F). Both the parental and knockout cells exhibited dome-like morphology; however, GFP expression was observed only in typical naïve clones from parental cells, not in the KLF17 knockout cells (Fig. 5G). This finding provided additional confirmation of the KLF17 knockout.

To rapidly assess the effect of KLF17 knockout during naïve programming, we characterized the cells by measuring the expression of naïve pluripotency-related genes in the 4CL medium from both parental and knockout lines (Fig. 5H). After 26 days of programming, both cell lines demonstrated significant increases in NANOG, DPPA3, DPPA5, and DNMT3L levels, though the knockout cells had lower levels of NANOG, DPPA3, and DNMT3L compared to parental cells. We also assessed KLF17 expression in both cell types during programming. The KLF17 knockout line showed a significant decrease in KLF17 mRNA, suggesting that the truncation of the KLF17 coding sequence caused by frameshift knockouts might lead to transcript degradation via nonsense-mediated mRNA decay (NMD)^30^. These findings are consistent with previous studies showing that KLF17 promotes human naïve pluripotency but is not essential for its establishment^31^.

## Discussion

Genome editing of silent genes in hPSCs does not impact the parental cells at the undifferentiated stage, as these targeted genes are not expressed. The phenotypic effects of genome editing only become evident when the genes are activated during differentiation. This creates a robust platform for studying targeted genes in the context of development or disease. However, the fact that these genes are silent in hPSCs presents a challenge in verifying the desired edits. Final verification of knock-in and knockout events typically relies on gene expression observed upon differentiation. Targeted PCR and Sanger sequencing of the genomic locus are essential for confirming gene knock-ins and knockouts in undifferentiated hPSCs. However, for knock-ins, some mutations might escape detection due to the limited length of PCR products and the resolution of Sanger sequencing. In the case of knockouts, it is crucial to assess the protein level of the gene, as some frameshift mutations might not fully eliminate the gene’s function, or wild-type cells may contaminate the knockout cells during single-cell clone generation.

In this study, we systematically applied CRISPRa, a technology for epigenetic activation of endogenous genes, for the effective verification of gene knock-ins and knockouts targeting silent genes in hPSCs. Previous studies have shown that the potency of different CRISPRa systems varies between cell lines. For example, SAM (Synergistic Activation Mediator) has shown more pronounced gene activation in HeLa cells, while Suntag and VPR were more effective in U-2 OS and MCF7 cells^32^. The SPH system was developed to yield higher activation of neural genes than SAM, Suntag, or VPR in both 293T and N2a cells^23^. In hPSCs, dCas9 fused with histone acetyltransferase p300 (dCas9-p300) showed limited effectiveness, while dCas9-VPR demonstrated success in most genomic contexts^33^. Despite the importance of selecting an appropriate CRISPRa tool for hPSC research, no comprehensive evaluation of different CRISPRa potencies in hPSCs has been available. Therefore, we compared three potent and widely used CRISPRa systems: VPR, SPH, and SAM, and found that SAM performed significantly better, achieving more robust gene activation than VPR and SPH within 48 hours across seven silent genes.

Moreover, several studies have indicated a combinatory effect of gene activation by CRISPRa with DNA demethylation for hypermethylated genes^34–36^. In our experiments, we utilized components of the SAM system, including the MS2 coat protein fused with three activator domains (VP64, P65, and HSF1) and an sgRNA-MS2 aptamer, along with dCas9 fused with TET1 at its N-terminal linked with an XTEN linker (TETv4). The TETv4 construct has been reported to enhance gene activation significantly and was applied effectively to reverse gene silencing mediated by CRISPRoff^37^. In our tests, the activation of KLF17 and HOPX genes was significantly enhanced by the inclusion of TET1 (Fig.2). Demethylation effects by TET1 were observed around the targeting region on the promoters of these two genes, particularly for KLF17, where most CpG sites are over 90% methylated in parental hPSC cells. Our data showed SAM-TET1 is the most effective tool to activate the gene loci with hypermethylation.

Based on our results demonstrating rapid activation of silent genes in hPSCs by CRISPRa, we propose a workflow for verifying different genome edits at silenced gene loci in hPSCs. We showed how to apply CRISPRa for the verification of three representative engineered hPSC lines: reporter hPSC lines, degron knock-in lines, and gene knockout lines at the undifferentiated hPSC stage. The highest gene activation was achieved around 48 hours after nucleofection (Fig. 4B, C), with gene expression gradually decreasing due to the degradation of plasmids delivered by transient transfection. Based on this observation, the verification time point was set to 48 hours across all experiments. The mRNA levels induced by SAM or SAM-TET1 ranged from over 70-fold to over 15,000-fold across the seven cases we tested, indicating robust gene activation (Fig.1D, 2F, 2G). The level of genes induced with this approach was higher than in differentiated cells. For example, SOX10 gene expression increased over 350-fold with SAM, compared to a 20-35-fold increase in SOX10-expressing neural crest cells relative to undifferentiated hPSCs based on RNA-seq data^38^ (Fig. 1D). In the case of KLF17 activation, SAM-TET1 induced a 9,000-fold increase in KLF17 expression, compared to a 7,000-fold increase in naïve cells expressing KLF17 (Fig.1F, 5H). These data indicate that this approach is sensitive enough to detect both gene knock-ins and knockouts. This also underscores the importance of careful sgRNA design to ensure robust gene activation.

Beyond reporter gene knock-ins and endogenous gene knockouts, we also demonstrated the functional assay of the dTAG system targeting endogenous loci, which includes temporal and reversible control of targeted protein degradation. The required time for protein degradation and its reversibility might differ in real cases where the endogenous gene is expressed in specific cell types derived from hPSCs. This discrepancy is due to the varying expression levels of the targeted protein and the duration of targeted gene activation. Nonetheless, functional evaluation in undifferentiated hPSCs with CRISPRa accurately reflects the selective control of the protein of interest by degron tags. This evaluation system can also be used to assess different chemical degraders targeting the same degron tag or to compare the efficacy of different degron systems targeting the same endogenous gene. In cases where longer gene activation is required for testing the functional assay of degrons, alternative delivery systems, such as lentivirus infection, can be considered.

## Methods

### Cell culture

All the hPSC lines are cultured on Matrigel (Fisher Scientific 08-774-552) in Stemflex Medium (Thermo Fisher A3349401) at 37 °C with 5% CO_2_. Fresh medium is replaced daily. Cells were passaged every 2-3 days with 1:4-1:6 ratio by incubation cells with 0.5 mM EDTA (Fisher Scientific MT-46034CI) for 5 min. All the hPSC lines were tested negative for mycoplasma contamination.

### Plasmid construction

For VPR activation, the dCas9-VPR plasmid (Addgene #114195), which expresses dCas9 fused with VP64, P65, and Rta, was used. For SPH activation, dCas9 fused with 10 copies of GCN4 (10xGCN4, Addgene #107310) and scFv-p65-HSF1 were cloned from the scFv-p65-HSF1-T2A-EGFP plasmid (Addgene #107311) by removing the EGFP from the plasmid. For SAM activation, the dCas9-VP64 plasmid (Addgene #61425) and MCP-P65-HSF1 plasmid (Addgene #89308) were used. For the SAM-TET1 system, the catalytic domain of TET1 (TET1CD) followed by the XTEN80 linker was amplified from the TETv4 plasmid (Addgene #176983) and cloned into the N-terminus of dCas9 under the EF1a promoter. The VP64 fragment was cloned into the MCP-P65-HSF1 plasmid to generate the MCP-VP64-P65-HSF1 plasmid. To clone sgRNA plasmids, the target sequence was cloned into the regular sgRNA backbone (Addgene #52963) for VPR and Suntag activation or into the MS2-sgRNA backbone (Addgene #73797) for SAM activation via BsmBI sites. Target sequence for CRISPRa were listed in Supplementary Table1.

For all donor plasmids used in HDR-mediated knock-in in hPSCs, plasmids containing 400-500 bp of left and right homology arms flanking the stop codon of the target gene were first cloned into the pUC57 vector. A P2A-reporter cassette followed by a PGK-Puro cassette flanked by loxP sites was cloned between the two homology arms using restriction enzyme sites. For the dTAG knock-in donor, the FKBP(F36V)-mCherry cassette was cloned into SOX10-GFP donor plasmids to replace the P2A-GFP. The sgRNA targeting sites for knock-in and knockout experiments were cloned into the px330 plasmid (Addgene #42230). Target sequence for sgRNAs were listed in Supplementary Table1.

### hPSC lines

Human embryonic stem cell lines H1 and H9 were purchased from the WiCell Institute. All hPSC reporter lines were generated through Cas9-mediated HDR as previously described^5^. The TBXT-GFP^5^, NGN2-GFP, and KLF17-GFP reporter lines were derived from H1 cells, while the other reporter lines, including MAP2-GFP, HOPX-GFP^26^, SOX10-GFP^24^, SOX10-dTAG-mCherry, OLIG2-tdTomato, and the dual reporter SOX10-GFP/OLIG2-tdTomato, were derived from H9 cells.

For the knock-in lines, 1 μg of the px330-sgRNA plasmid, or a pair of px330-sgRNA plasmids (1 μg each), and 1 μg of the donor plasmid were co-electroporated into the hPSCs. Cells were treated with 1 μg/ml puromycin for 72 hours, beginning 72 hours after electroporation. To generate knockout lines, 1 μg of the px330-sgRNA plasmid and a PGK-puro plasmid were co-transfected into H1 cells to knock out SOX10, and into H1-KLF17-GFP cells to knock out KLF17, via electroporation. Cells were treated with 0.5 μg/ml puromycin for 24 hours, 24 hours after electroporation. Puromycin-selected cells were then re-plated into a 96-well plate for single-cell clone generation and genotyped by Sanger sequencing.

### Electroporation in hPSCs

Cells were dissociated using Accutase (Innovative Cell Tech. AT104) and electroporated with plasmids for gene activation using P3 Primary Cell 4D-Nucleofector X Kit (Lonza V4XP-3032). The reactions were performed using the “CB-150” program on the Lonza 4D-Nucleofector X Unit. The cells from one reaction were resuspended with Stemflex supplemented with 10 μM Y-27632 (Stemcell Technologies) and replated into one well of a Matrigel coated 48-well plate.

### FACS analysis of reporter gene

Reporter lines were dissociated with Accutase, resuspended in cold PBS containing 0.5% BSA and filtered through a cell strainer with 35μm sized mesh (Fisher Scientific 352235) to remove clumps. Reporter gene expressing cells were analyzed using a BD FACSAria III (BD Bioscience). Data analysis was conducted with FCS Express software (version 7.18.0025, DeNovo Software) with. Gating strategy is showed in Supplementary Fig. 8.

### Assessment of Cell viability

The viability of the cells treated with different CRISPRa systems was determined by Acridine orange (AO) /propidium iodide (PI) staining 48 hours after electroporation. Live cells stained with AO in green and dead cells stained with PI presented in red and the total cell number were calculated with cell counter (Nexcelom Bioscience).

### Quantitative realtime PCR

For the comparison of gene activation in hPSCs with different CRISPRa systems, cells were isolated 48 hours after electroporation for RNA extraction. To evaluate the expression of naïve hPSC markers, H1-KLF17-GFP cells and the corresponding KLF17 knockout cells were collected on Day 26 of naïve programming. RNA was extracted using the RNeasy Mini Kit (Qiagen, 74104) and subsequently synthesized into cDNA with the SuperScript VILO Master Mix (Invitrogen, 11755050). Quantitative real-time PCR was performed with PowerUp™ SYBR™ Green Master Mix (Applied Biosystems™) on a QuantStudio 5 (Applied Biosystems). Primers for the targeted genes are listed in Supplementary Table 2.

### Targeted bisulfite sequencing

Genomic DNA was extracted from the cells 48 hours after electroporation and subjected to targeted bisulfite sequencing. Primers were specifically designed to target CpG sites around the sgRNA targeting regions. The DNA samples were bisulfite-converted using the EZ DNA Methylation-Lightning™ Kit (Zymo Research). The targeted regions were then amplified via PCR using the designed primers, and the resulting amplicons were pooled for barcoding. Samples were prepared using a MiSeq V2 300bp Reagent Kit (Illumina) for paired-end sequencing. Low-quality nucleotides and adapter sequences were trimmed during quality control analysis using Trim Galore (https://www.bioinformatics.babraham.ac.uk/projects/trim_galore/). FastQC (https://www.bioinformatics.babraham.ac.uk/projects/fastqc/) was used to evaluate the sequencing quality of both the raw and trimmed reads. Sequence reads were aligned back to the reference genome using Bismark (http://www.bioinformatics.babraham.ac.uk/projects/bismark/). The methylation level of each sampled cytosine was estimated by dividing the number of reads reporting a C by the total number of reads reporting a C or T.

### Flow cytometry

Cell populations with reporter gene activation at the undifferentiated hPSC stage were evaluated using FACS 48 hours after electroporation with CRISPRa. Cells were dissociated into single cells using Accutase (Innovative Cell Tech. AT104) and resuspended in cold PBS containing 0.5% BSA. Reporter gene-expressing cells were then analyzed using a BD FACSAria III (BD Bioscience) and the data were processed with FCS Express software (version 7.18.0025, DeNovo Software).

### Generation of oligodendrocyte progenitors with H9-SOX10-GFP/OLIG2-tdTomato dual reporter cells

H9-SOX10-GFP/OLIG2-tdTomato dual reporter cells were first converted to neural stem cells (NSCs) by dual-SMAD inhibition, and then specified to the ventral spinal cord domain by the caudalization activity of CHIR, a potent WNT-pathway activator, and retinoic acid (RA). Within the developing spinal cord, the neural cells pool is restricted to the pMN domain that, at first, generates motor neurons, and later oligodendrocytes. The main signaling pathways involved in maintaining the pMN domain are sonic hedgehog and retinoic acid, which are kept high in the protocol from this point on. As cells are specified to the pMN, they start expressing OLIG2, which can either direct their fate into neurons or oligodendrocytes. By providing a glia competency cocktail that includes PDGFRa ligand, we push OLIG2+ cells to become glia competent expressing NFIA and SOX9. Finally, we provide a terminal differentiation cocktail which allows the cells to differentiate into oligodendrocyte progenitors (OPCs) expressing nuclear SOX10. As SOX10+ OPCs mature, they will start acquiring a unique combinations of surface markers such as O4 and O1 until becoming fully mature MBP+ oligodendrocytes.

### Naïve programming

Naïve PSCs were generated using 4CL medium as previously described^29^. Primed H1-KLF17-GFP cells and the KLF17 knockout line were dissociated using Accutase (Innovative Cell Tech. AT104). Single cells were plated at a density of 1,000 to 1,500 cells/cm² on irradiated CF1 mouse embryonic fibroblasts (Gibco) in Stemflex Medium (Thermo Fisher A3349401) supplemented with 10 μM Y-27632 (Stemcell Technologies). After 24 hours, the culture medium was replaced with 4CL medium, composed of a 1:1 mix of Neurobasal medium (Gibco, 21103049) and Advanced DMEM/F12 (Gibco, 12634028), supplemented with N2 (Gibco, 17502048) and B27 (Gibco, 17504044), sodium pyruvate (Corning, 25000CL), non-essential amino acids (Corning, 25025CL), GlutaMAX (Gibco, 35050061), penicillin-streptomycin (HyClone, SV30010), 10 nM DZNep (Selleck, S7120), 5 nM TSA (Vetec, V900931), 1 µM PD0325901 (Axon, 1408), 5 µM IWR-1 (Sigma, I0161), 20 ng/ml human LIF (Peprotech, 300-05), 20 ng/ml activin A (Peprotech, 120-14E), 50 µg/ml l-ascorbic acid (Sigma, A8960), and 0.2% (v/v) Matrigel. Cells were fed with fresh 4CL medium daily and passaged with Accutase at a 1:5 to 1:8 ratio every 3 to 4 days.

### Western blot

SOX10 was activated in both parental H1 cells and SOX10 knockout cells using SAM-TET1 for 48 hours. Cells were harvested and lysed in RIPA buffer (Thermo Fisher Scientific) supplemented with 1× HALT protease inhibitor cocktail (Thermo Fisher Scientific) and 5 mM EDTA. Protein concentrations were determined using a BCA assay (Pierce), and equal amounts of protein from different samples were loaded onto a 4–15% precast gel (Bio-Rad). SOX10 was detected using a mouse anti-SOX10 antibody (1:1000, sc-271163, Santa Cruz Biotechnology). α-Tubulin was used as a protein loading control (1:2000, #2144, Cell Signaling Technology).

## Acknowledgments

This work was supported by NIH/NCI Cancer Center Support Grant P30 CA008748 from Memorial Sloan Kettering. The work was also supported by a core facility grant of the Starr Foundation through the Tri-Institutional Stem Cell Initiative and by the Contract C029153 from the New York State’s stem cell funding agency (NYSTEM).

## Data availability statement

The plasmids generated in this study have been deposited and pending in Addgene. All other data supporting the findings of this study are available from the corresponding authors upon reasonable request.

## Code availability statement

The Benchling CRISPR Design website was used for design sgRNA or pegRNA spacer and is available at (https://www.benchling.com/#).

## Author contributions

Y.W., T.Z., L.S. conceived the project and designed the experiments. Y.W., A.Z., A.E., M.S., T.Z. performed experiments and analyzed data. Y.W., A.E., T.Z., L.S. wrote the manuscript.

---

**Fig. S1:**
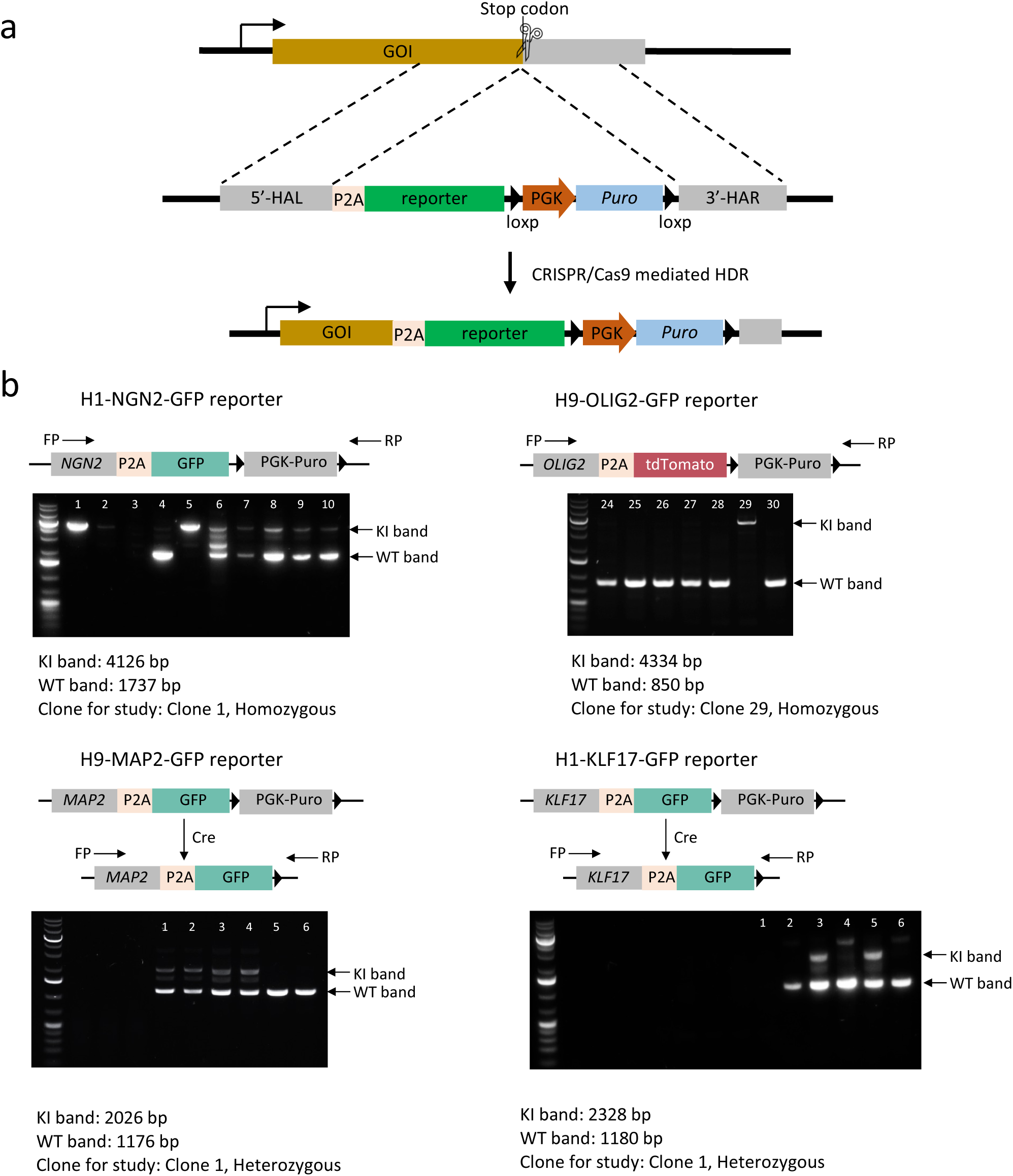
Reporter line generation using CRISPR/Cas9 mediated HDR in hPSCs. **a.** Schematic of knocking in of reporter gene in the gene of interest (GOI) using CRISPR Cas9. **b.** PCR verification of reporter gene knock-in at the *NGN2*, *OLIG2*, *MAP2*, *KLF17* and *HOPX* loci. Knock-in (KI) band and non-knock-in (WT) bands are indicated on the gel.

**Fig. S2:**
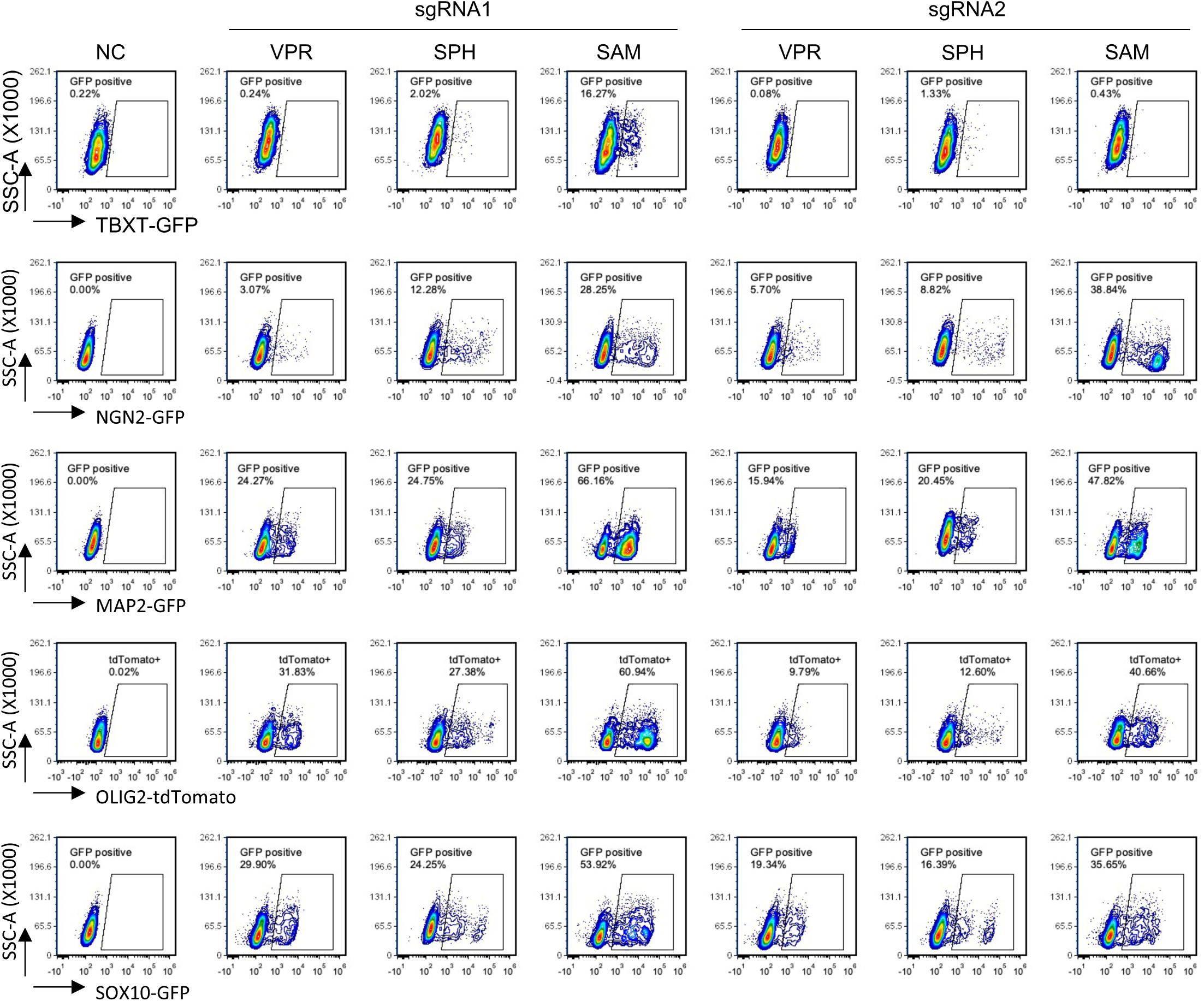
Representative FACS plots showing the reporter gene-expressing cells activated by different CRISPRa systems in the hPSC reporter lines 48 hours after electroporation.

**Fig. S3:**
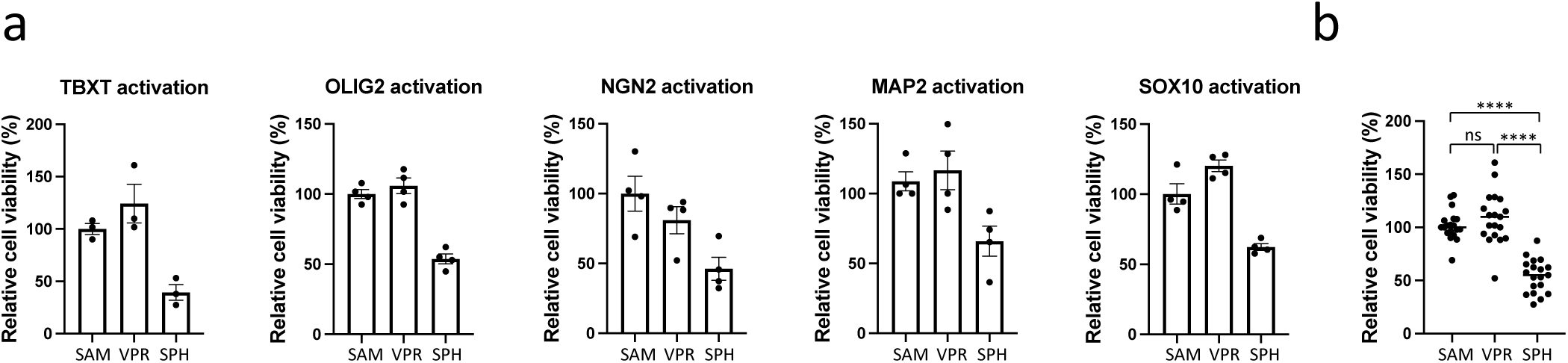
Viability of the cells with SAM, VPR or SPH treatment. **a.** Cell viability was determined by AO/PI staining 48 hours after TBXT, OLIG2, NGN2, MAP2 and SOX10 activation using SAM, VPR and SPH CRISPRa systems. The data are normalized to the viability in the SAM-treated cells. Data represent three independent experiments for TBXT activation and four independent experiments for the other four gene activations. **b.** Comparison of cell viability with the three CRISPRa systems targeting five loci from a. The center line shows the medians of all the data points from the five target genes. *p* values were calculated by one-way ANOVA with Tukey’s multiple comparison test (ns: non-significant, \*\*\*\**p*<0.0001).

**Fig. S4.**
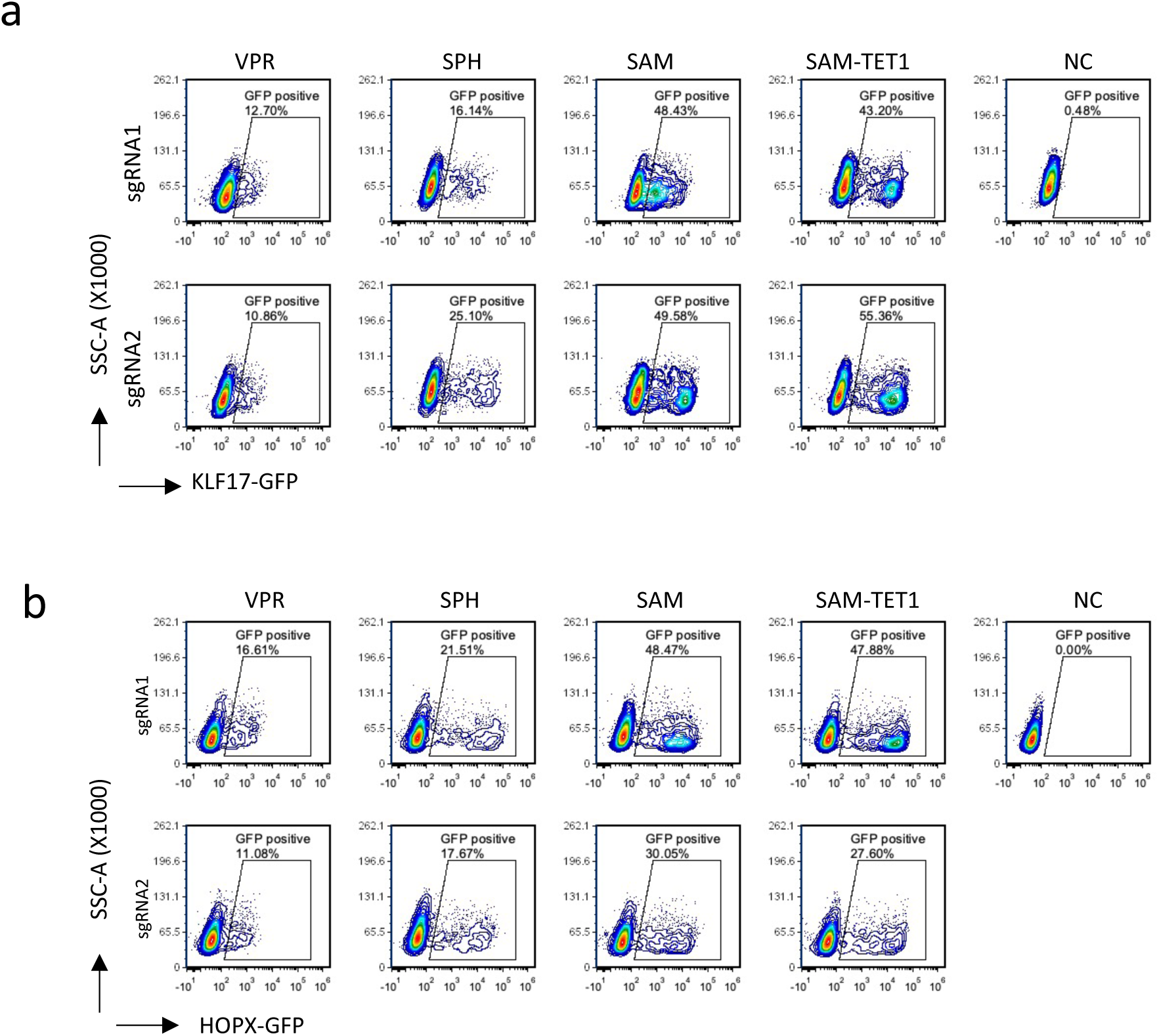
Representative FACS plots showing the GFP reporter gene-positive cells induced by VPR, SPH, SAM and SAM-TET1. **a.** GFP-positive cells in the H1-KLF17-GFP cells with KLF17 activation by the indicated systems for 48 hours. **b.** GFP-positive cells in the H9-HOPX-GFP cells with KLF17 activation by the indicated systems for 48 hours. *p* values were calculated by one-way ANOVA with Šídák’s multiple comparisons test as a post-hoc test (ns: non-significant, \*\**p*<0.01, \*\*\**p*<0.001, \*\*\*\**p*<0.0001).

**Fig. S5.**
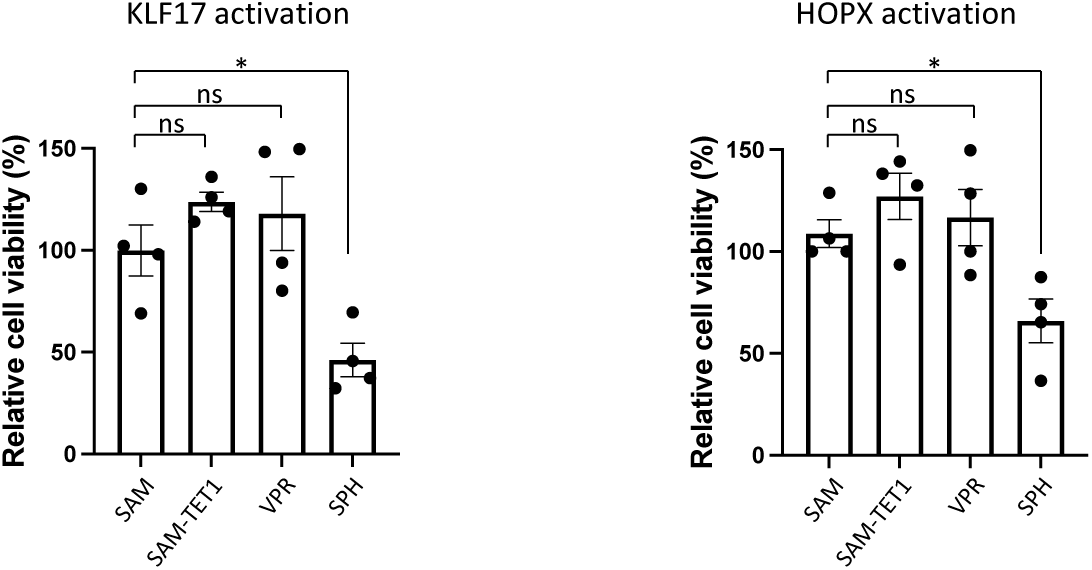
Viability of the cells after activation of KLF17 and HOPX with VPR, SPH, SAM and SAM-TET1 for 48 hours. Bars represent data from four independent experiments. *p* values were calculated by one-way ANOVA followed by Dunnett’s test (ns: non-significant, \**p*<0.05).

**Fig. S6:**
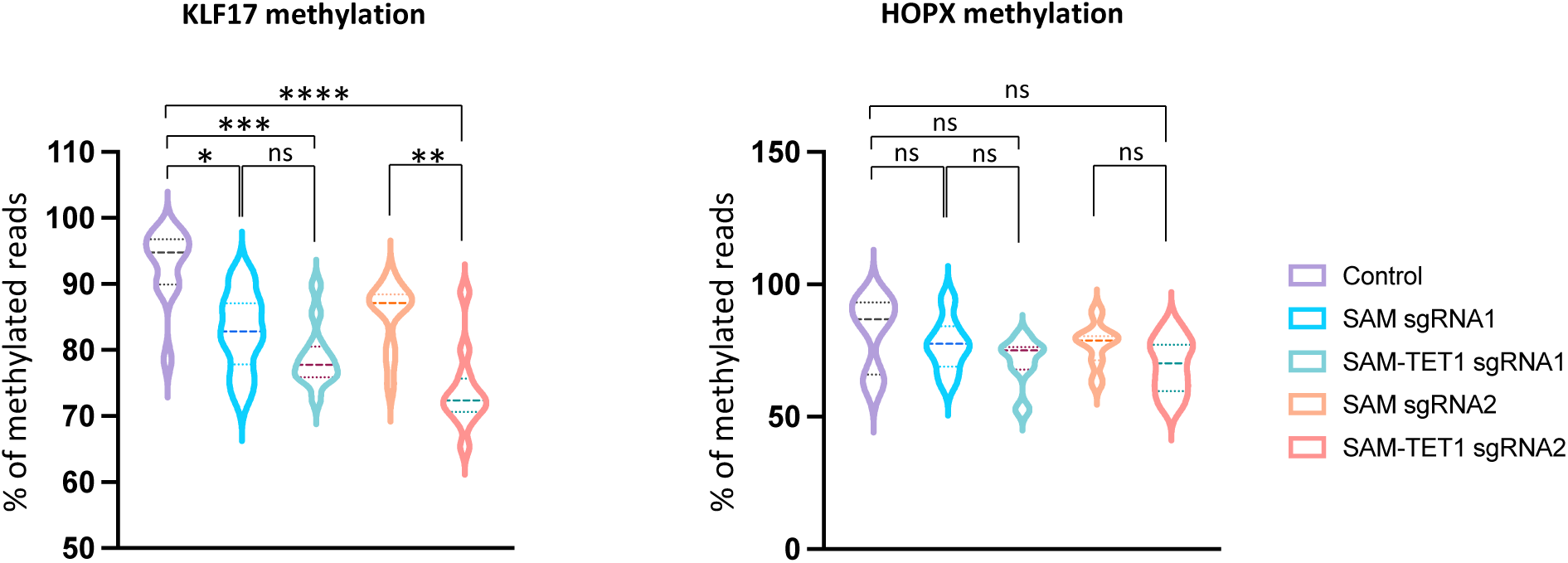
Comparison of the methylation ratio around sgRNA targeted regions of the KLF17 and HOPX genes by SAM and SAM-TET1. The statistical significance between the negative control, SAM and SAM-TET1 groups for the entire set of CpG sites from the data in Fig. 2I and 2K data was calculated using Kruskal-Wallis test *p*-values, followed by Dunn’s multiple comparison test (ns: non-significant, *p<0.05, \*\**p*<0.01, \*\*\**p*<0.001, \*\*\*\**p*<0.0001).

**Fig. S7.**
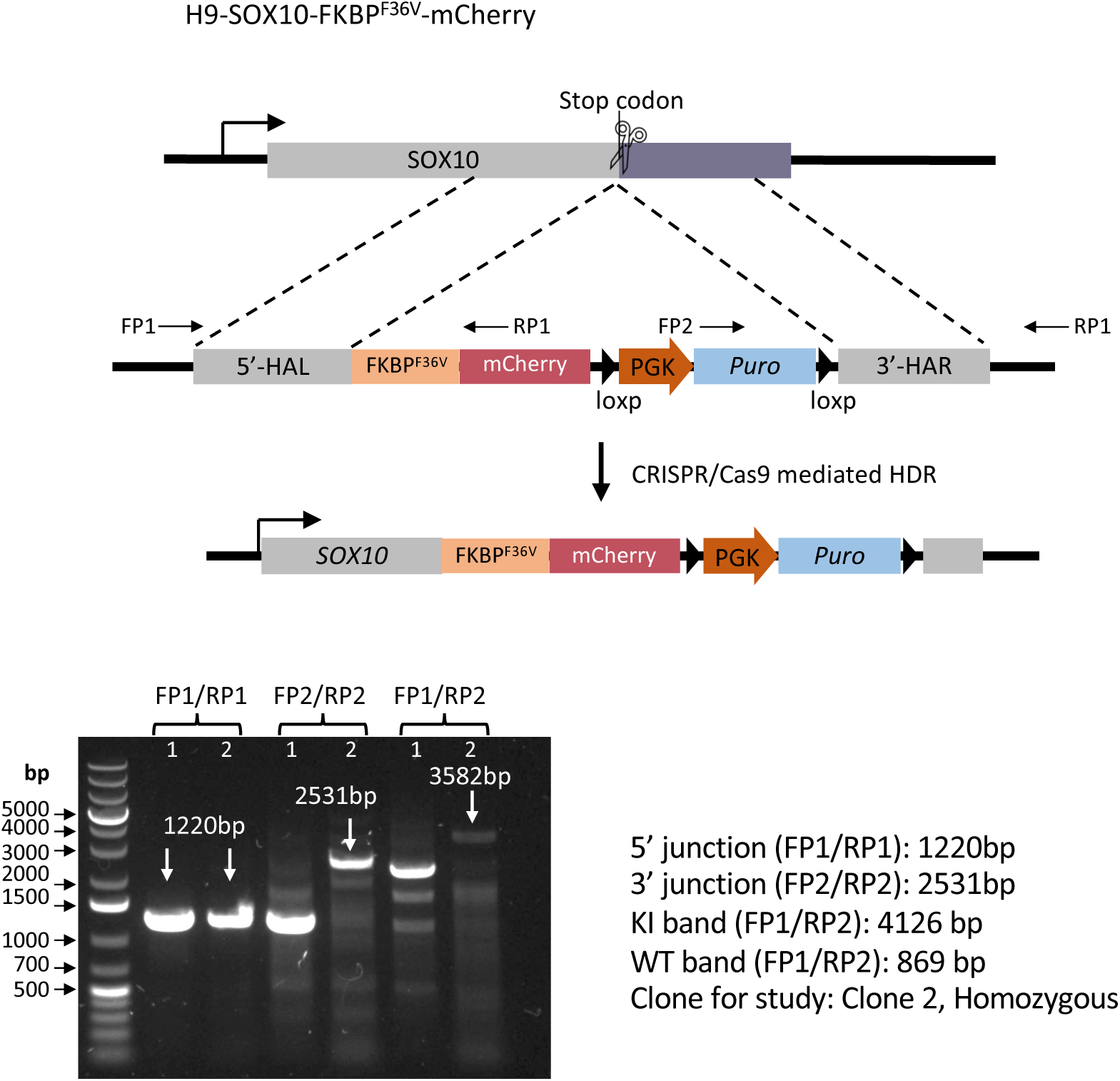
Generation of dTAG knock-in line at the SOX10 locus. a. Schematic of knock-in of FKBP^F36V^-mCherry sequence in-frame with SOX10 using CRISPR/Cas9-mediated HDR. b. PCR verification of dTAG gene knock-in.

**Fig. S8:**
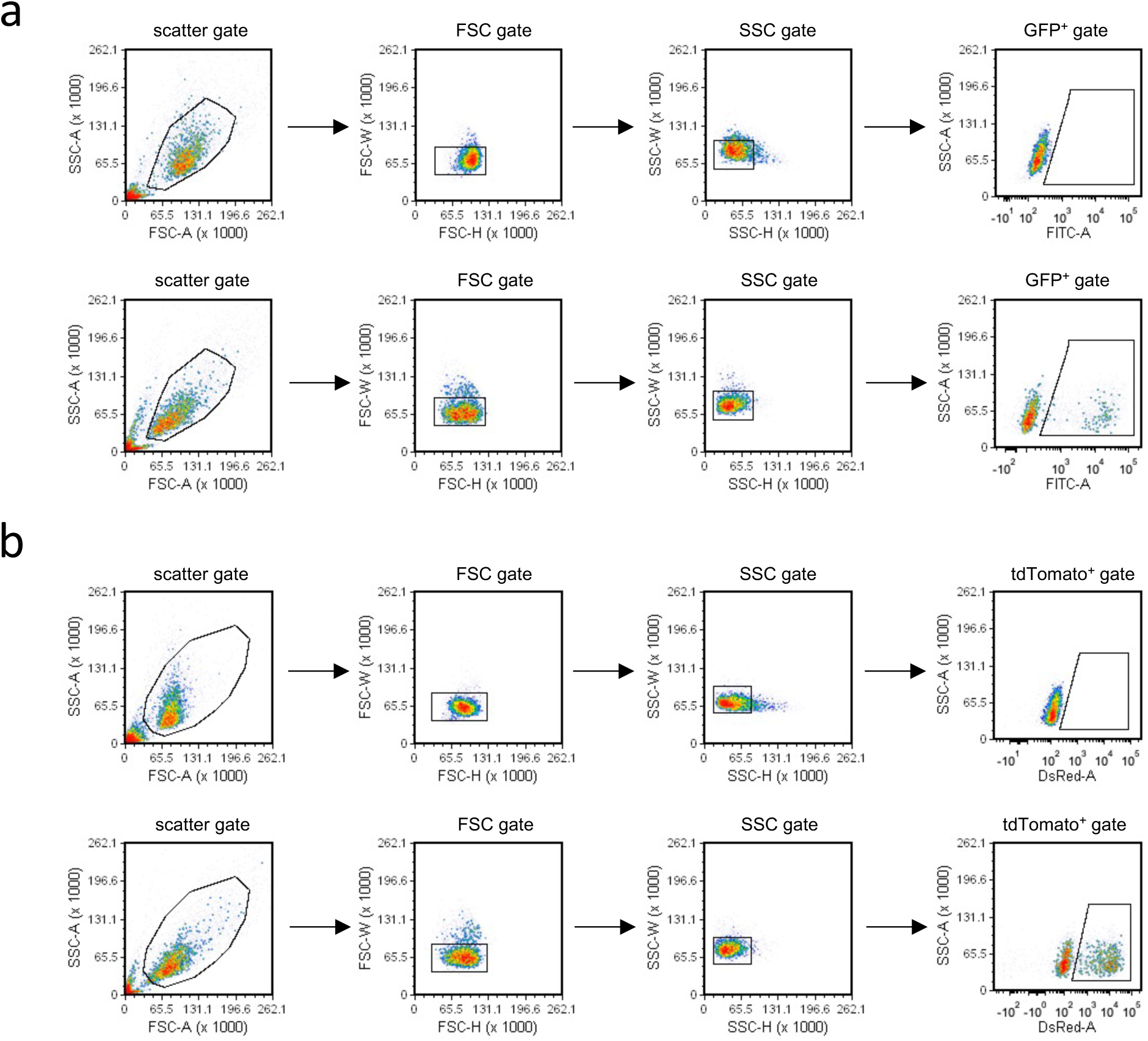
FACS gating examples for the detection of GFP (a) and tdTomato (b) positive cells. Cells were initially gated on population using FSC-A/SSC-A (scatter gate). Single cells were gated using FSC-W/FSC-H (FSC gate) and SSC-W/SSC-H (SSC gate). The GFP positive cells and tdTomato positive cell population were determined relative to the negative control.

## References

1 Ben Jehuda, R., Shemer, Y. & Binah, O. Genome Editing in Induced Pluripotent Stem Cells using CRISPR/Cas9. Stem Cell Rev Rep 14, 323–336 (2018). 10.1007/s12015-018-9811-3

2 Hendriks, W. T., Warren, C. R. & Cowan, C. A. Genome Editing in Human Pluripotent Stem Cells: Approaches, Pitfalls, and Solutions. Cell stem cell 18, 53–65 (2016). 10.1016/j.stem.2015.12.002

3 Grskovic, M., Javaherian, A., Strulovici, B. & Daley, G. Q. Induced pluripotent stem cells- -opportunities for disease modelling and drug discovery. Nat Rev Drug Discov 10, 915–929 (2011). 10.1038/nrd3577

4 Soldner, F. & Jaenisch, R. Stem Cells, Genome Editing, and the Path to Translational Medicine. Cell 175, 615–632 (2018). 10.1016/j.cell.2018.09.010

5. Zhong, A., Li, M. & Zhou, T. Protocol for the Generation of Human Pluripotent Reporter Cell Lines Using CRISPR/Cas9. STAR protocols 1 (2020). 10.1016/j.xpro.2020.100052

6 Dewari, P. S. et al. An efficient and scalable pipeline for epitope tagging in mammalian stem cells using Cas9 ribonucleoprotein. Elife 7 (2018). 10.7554/eLife.35069

7 Borowiak, M. et al. Small molecules efficiently direct endodermal differentiation of mouse and human embryonic stem cells. Cell Stem Cell 4, 348–358 (2009). 10.1016/j.stem.2009.01.014

8 Longmire, T. A. et al. Efficient derivation of purified lung and thyroid progenitors from embryonic stem cells. Cell Stem Cell 10, 398–411 (2012). 10.1016/j.stem.2012.01.019

9 Zhu, W. et al. Precisely controlling endogenous protein dosage in hPSCs and derivatives to model FOXG1 syndrome. Nat Commun 10, 928 (2019). 10.1038/s41467-019-08841-7

10 Nabet, B. et al. The dTAG system for immediate and target-specific protein degradation. Nat Chem Biol 14, 431–441 (2018). 10.1038/s41589-018-0021-8

11 Li, S., Prasanna, X., Salo, V. T., Vattulainen, I. & Ikonen, E. An efficient auxin-inducible degron system with low basal degradation in human cells. Nat Methods 16, 866–869 (2019). 10.1038/s41592-019-0512-x

12 Xu, H. et al. Targeted Disruption of HLA Genes via CRISPR-Cas9 Generates iPSCs with Enhanced Immune Compatibility. Cell Stem Cell 24, 566–578 e567 (2019). 10.1016/j.stem.2019.02.005

13 Li, C. et al. Single-cell brain organoid screening identifies developmental defects in autism. Nature 621, 373–380 (2023). 10.1038/s41586-023-06473-y

14 Ding, Q., Regan, S. N., Xia, Y., Oostrom, L. A., Cowan, C. A. & Musunuru, K. Enhanced efficiency of human pluripotent stem cell genome editing through replacing TALENs with CRISPRs. Cell Stem Cell 12, 393–394 (2013). 10.1016/j.stem.2013.03.006

15 Maeder, M. L., Linder, S. J., Cascio, V. M., Fu, Y., Ho, Q. H. & Joung, J. K. CRISPR RNA-guided activation of endogenous human genes. Nat Methods 10, 977–979 (2013). 10.1038/nmeth.2598

16 Cheng, A. W. et al. Multiplexed activation of endogenous genes by CRISPR-on, an RNA-guided transcriptional activator system. Cell Res 23, 1163–1171 (2013). 10.1038/cr.2013.122

17 Weltner, J. et al. Human pluripotent reprogramming with CRISPR activators. Nat Commun 9, 2643 (2018). 10.1038/s41467-018-05067-x

18 Liu, P., Chen, M., Liu, Y., Qi, L. S. & Ding, S. CRISPR-Based Chromatin Remodeling of the Endogenous Oct4 or Sox2 Locus Enables Reprogramming to Pluripotency. Cell Stem Cell 22, 252–261 e254 (2018). 10.1016/j.stem.2017.12.001

19 Liu, Y. et al. CRISPR Activation Screens Systematically Identify Factors that Drive Neuronal Fate and Reprogramming. Cell Stem Cell 23, 758–771 e758 (2018). 10.1016/j.stem.2018.09.003

20 Kwon, J. B., Vankara, A., Ettyreddy, A. R., Bohning, J. D. & Gersbach, C. A. Myogenic Progenitor Cell Lineage Specification by CRISPR/Cas9-Based Transcriptional Activators. Stem Cell Reports 14, 755–769 (2020). 10.1016/j.stemcr.2020.03.026

21 Chavez, A. et al. Highly efficient Cas9-mediated transcriptional programming. Nat Methods 12, 326–328 (2015). 10.1038/nmeth.3312

22 Konermann, S. et al. Genome-scale transcriptional activation by an engineered CRISPR-Cas9 complex. Nature 517, 583–588 (2015). 10.1038/nature14136

23 Zhou, H. et al. In vivo simultaneous transcriptional activation of multiple genes in the brain using CRISPR-dCas9-activator transgenic mice. Nat Neurosci 21, 440–446 (2018). 10.1038/s41593-017-0060-6

24 Chambers, S. M. et al. Combined small-molecule inhibition accelerates developmental timing and converts human pluripotent stem cells into nociceptors. Nat Biotechnol 30, 715–720 (2012). 10.1038/nbt.2249

25 Morita, S. et al. Targeted DNA demethylation in vivo using dCas9-peptide repeat and scFv-TET1 catalytic domain fusions. Nat Biotechnol 34, 1060–1065 (2016). 10.1038/nbt.3658

26 Walsh, R. M. et al. Generation of human cerebral organoids with a structured outer subventricular zone. Cell Rep 43, 114031 (2024). 10.1016/j.celrep.2024.114031

27 Guo, G. et al. Naive Pluripotent Stem Cells Derived Directly from Isolated Cells of the Human Inner Cell Mass. Stem Cell Reports 6, 437–446 (2016). 10.1016/j.stemcr.2016.02.005

28 Blakeley, P. et al. Defining the three cell lineages of the human blastocyst by single-cell RNA-seq. Development 142, 3613 (2015). 10.1242/dev.131235

29 Mazid, M. A. et al. Rolling back human pluripotent stem cells to an eight-cell embryo-like stage. Nature 605, 315–324 (2022). 10.1038/s41586-022-04625-0

30 Tuladhar, R. et al. CRISPR-Cas9-based mutagenesis frequently provokes on-target mRNA misregulation. Nat Commun 10, 4056 (2019). 10.1038/s41467-019-12028-5

31 Lea, R. A. et al. KLF17 promotes human naive pluripotency but is not required for its establishment. Development 148 (2021). 10.1242/dev.199378

32 Chavez, A. et al. Comparison of Cas9 activators in multiple species. Nat Methods 13, 563–567 (2016). 10.1038/nmeth.3871

33 Wu, Q. et al. Massively parallel characterization of CRISPR activator efficacy in human induced pluripotent stem cells and neurons. Mol Cell 83, 1125–1139 e1128 (2023). 10.1016/j.molcel.2023.02.011

34 Morita, S., Horii, T., Kimura, M. & Hatada, I. Synergistic Upregulation of Target Genes by TET1 and VP64 in the dCas9-SunTag Platform. Int J Mol Sci 21 (2020). 10.3390/ijms21051574

35 Halmai, J. et al. Artificial escape from XCI by DNA methylation editing of the CDKL5 gene. Nucleic Acids Res 48, 2372–2387 (2020). 10.1093/nar/gkz1214

36 Chan, W. F. et al. Activation of stably silenced genes by recruitment of a synthetic de-methylating module. Nat Commun 13, 5582 (2022). 10.1038/s41467-022-33181-4

37 Nunez, J. K. et al. Genome-wide programmable transcriptional memory by CRISPR-based epigenome editing. Cell 184, 2503–2519 e2517 (2021). 10.1016/j.cell.2021.03.025

38 Fan, Y. et al. hPSC-derived sacral neural crest enables rescue in a severe model of Hirschsprung’s disease. Cell stem cell 30, 264–282 e269 (2023). 10.1016/j.stem.2023.02.003

